# Task modulation of spatiotemporal dynamics in semantic brain networks: an EEG/MEG study

**DOI:** 10.1101/2021.06.28.450126

**Authors:** Setareh Rahimi, Seyedeh-Rezvan Farahibozorg, Rebecca Jackson, Olaf Hauk

## Abstract

How does brain activity in distributed semantic brain networks evolve over time, and how do these regions interact to retrieve the meaning of words? We compared spatiotemporal brain dynamics between visual lexical and semantic decision tasks (LD and SD), analysing whole-cortex evoked responses and spectral functional connectivity (coherence) in source-estimated electroencephalography and magnetoencephalography (EEG and MEG) recordings. Our evoked analysis revealed generally larger activation for SD compared to LD, starting in primary visual area (PVA) and angular gyrus (AG), followed by left posterior temporal cortex (PTC) and left anterior temporal lobe (ATL). The earliest activation effects in ATL were significantly left-lateralised. Our functional connectivity results showed significant connectivity between left and right ATLs and PTC and right ATL in an early time window, as well as between left ATL and IFG in a later time window. The connectivity of AG was comparatively sparse. We quantified the limited spatial resolution of our source estimates via a leakage index for careful interpretation of our results. Our findings suggest that semantic task demands modulate visual and attentional processes early-on, followed by modulation of multimodal semantic information retrieval in ATLs and then control regions (PTC and IFG) in order to extract task-relevant semantic features for response selection. Whilst our evoked analysis suggests a dominance of left ATL for semantic processing, our functional connectivity analysis also revealed significant involvement of right ATL in the more demanding semantic task. Our findings demonstrate the complementarity of evoked and functional connectivity analysis, as well as the importance of dynamic information for both types of analyses.

**Highlights:**

1. Semantic task demands affect activity and connectivity at different processing stages
2. Earliest task modulations occurred in posterior visual brain regions
3. ATL, PTC and IFG effects reflect task-relevant retrieval of multimodal information
4. ATL effects left-lateralised for activation but bilateral for functional connectivity
5. Dynamic evoked and connectivity data are essential to study semantic networks

## 1 Introduction

Semantics, or the representation and mental manipulation of our knowledge about objects, facts and people, is a crucial component of human cognition, underpinning all meaningful interactions with our environment and communication with others (Jefferies, 2013; Patterson et al., 2007). Our semantic system enables us to store, employ, manipulate, and generalise conceptual knowledge (Lambon Ralph et al., 2016). Learning and storing multimodal semantic representations are essential for successful semantic cognition, but they are not sufficient. The relevant information to deploy in any particular moment is context-sensitive and task-dependent, thus, we require semantic control to manipulate and shape the activation in the representation system (Jackson, 2021; Jefferies, 2013). However, task effects on brain dynamics during semantic processing are still largely unexplored.

The Controlled Semantic Cognition (CSC) framework proposes an interaction between control and representation regions in the brain, with semantic representation underpinned by a central semantic hub located in the anterior temporal lobes (ATL) (Lambon Ralph et al., 2016). Ample evidence for this proposal has been provided by studies on semantic dementia patients, who show specific semantic deficits following impairment of the anterior temporal lobes (Mion et al., 2010; Nestor et al., 2006), and fMRI and PET studies demonstrating ATL sensitivity to semantic stimulus and task manipulations (Crinion et al., 2003; Embleton et al., 2006; Mummery et al., 2000; Rogers et al., 2006; Tranel et al., 2005; Visser et al., 2012, 2010). Several studies have demonstrated similar effects in brain activity estimated from EEG or MEG data (Cope et al., 2020; Dhond et al., 2007; Farahibozorg et al., 2019; Marinkovic et al., 2014, 2003; Mollo et al., 2017), but the precise time course of semantic processing, as reflected in the brain activation or connectivity measures, has not been established yet. As a result, crucial evidence for the dynamic functional organisation of the semantic brain network is still missing, since temporal information is essential to disentangle effects that may occur at different stages of semantic processing, e.g., early semantic information retrieval, control processes in decision making, and later imagery or episodic memory processes (Hauk, 2016). Temporal information is also particularly important for the reliable estimation of brain connectivity, since brain areas may play different roles at different stages of processing, and therefore dynamically change their connectivity. Furthermore, there is evidence that activity in different brain networks, arguably corresponding to different brain functions, are reflected in different frequency bands of electrical brain signals, such as EEG and MEG (Siegel et al., 2012).

Whilst ATL has consistently been linked to semantic representation (Acosta-Cabronero et al., 2011; Binder et al., 2016; Martin, 2016; Pobric et al., 2007; Rogers et al., 2004), IFG and pMTG are specifically implicated in semantic control (Badre et al., 2005; Jackson, 2021; Jefferies, 2013; Jefferies and Lambon Ralph, 2006; Lambon Ralph et al., 2016; Noonan et al., 2013). The role of AG is less clear and has been suggested to involve semantic representation (Binder et al., 2009), control (Noonan et al., 2013) or episodic memory processes (Humphreys et al., 2015). Thus, semantic cognition is dependent on semantic representation in the ATL and sensory-specific regions, and control in IFG and pMTG, with a possible role for the AG. Few studies have investigated the interaction between semantic control and representation regions, and the connectivity and temporal dynamics of the corresponding brain regions are still not well understood (Jefferies, 2013; Lambon Ralph et al., 2016). Most previous studies of the semantic network and its connectivity have employed fMRI (Alam et al., 2021; Chiou et al., 2018; Chiou and Lambon Ralph, 2019; Humphreys et al., 2015; Jackson et al., 2016; Kuhnke et al., 2020) which despite its excellent spatial resolution, is limited in tracking any neural response faster than one second.

In the present study, we employed EEG/MEG to provide novel evidence of the timing of task modulations of activity in and functional connectivity among these regions. EEG and MEG are sensitive to semantic stimulus manipulations in different time windows, such as the N400 latency range (typically between 250 to 500ms) (Kutas and Federmeier, 2011; Lau et al., 2008) and earlier (Amsel et al., 2013; Hauk et al., 2012; Pulvermüller et al., 2009). Importantly, source estimation with MEG has revealed lexicosemantic effects in the anterior and middle temporal lobes (Dhond et al., 2007; Farahibozorg et al., 2019; Flick et al., 2018; Hauk et al., 2012; Lau et al., 2013; Mollo et al., 2017), inferior parietal cortex (Bemis and Pylkkänen, 2013; Farahibozorg et al., 2019; Lewis et al., 2015; Williams et al., 2017) and inferior frontal cortex (Schoffelen et al., 2017; Woodhead et al., 2014). Furthermore, semantic task manipulations have been reported to modulate EEG/MEG signals in early and late time windows (Chen et al., 2015, 2013) and in the frequency domain (Clarke et al., 2011; Lewis and Bastiaansen, 2015; Mollo et al., 2017).

In the present study, we investigated the effects of different semantic task demands on dynamic brain activity and spectral functional connectivity in the semantic brain networks. We used a whole-cortex approach initially, but also focused on prominent regions-of-interest (ROIs) that have previously been implicated in semantic representation and control, as described above: ATL, IFG, pMTG, and AG. Most previous studies have found the semantic brain network to be left-lateralised (Binder et al., 2009), yet, a notable exception is the ATL, for which a graded lateralisation has been reported depending on stimulus and task features (Lambon Ralph et al., 2010; Marinkovic et al., 2003; Olson et al., 2007; Patterson et al., 2007; Pobric et al., 2007; Rice et al., 2015b, 2015a; Visser et al., 2010). Thus, our ROIs will include both left and right ATLs to study the laterality of task effects in this region.

We contrasted brain dynamics between two visual word recognition tasks, namely lexical and semantic decisions on the identical word stimuli. In the lexical decision (LD) task, participants had to distinguish between words and pseudowords. This task only explicitly requires the classification of letters strings as existing words or not, and therefore does not explicitly demand the retrieval of much semantic information. However, the harder the distinction between the words and pseudowords, the more these decisions are affected by semantic variables, and lexical decision is compromised with impaired semantic representations (Evans et al., 2012; Patterson et al., 2006). This task is therefore suitable to evoke activity in the semantic network. We compared this task with a semantic decision (SD) task which explicitly required participants to retrieve specific semantic information about the words (such as “Is it something edible with a distinctive odour?”). This ‘task differences’ approach (Chen et al., 2015, 2013; Kuhnke et al., 2021, 2020) employs a high-level baseline, providing a powerful way to identify specific changes with greater semantic processing, such as the particular timing of differences. By presenting the same stimuli in two different tasks we can assess the effect of demanding semantic processing over and above the effect of presenting meaningful stimuli.

1. We used spectral coherence as a functional connectivity metric, which is sensitive to covariations of both phase and amplitude between two signals (Bastos and Schoffelen, 2016). We investigated the potential effect of source leakage in an explicit resolution analysis of our measurement configuration (Hauk et al., 2019). Specifically, we asked 1) how task modulation of semantic brain activity evolves across time, 2) how connectivity of putative semantic representation and control regions is affected by task demands over time, and 3) how task demands modulate the laterality and connectivity of left and right ATLs.

## 2 Materials and Methods

### 2.1 EEG/MEG experiment data acquisition

#### 2.1.1 Participants

26 healthy native adult English speakers (age 18-40) participated, 2 of whom were excluded due to problems with structural MRI scans. 3 were excluded due to inadequate behavioural response accuracies (less than 75% response accuracy) and 3 were excluded because of excessive movement artefacts. The excessive movement artefacts were determined based on: visual inspection by two authors, number of bad channels and number of bad epochs. Therefore, 18 participants (mean age 27.00±5.13, 12 female) entered the final analysis. A reduced version of the Oldfield handedness inventory (Oldfield 1971) was used, based on which a mean handedness laterality quotient of 89.84±0.2 was obtained. All participants had normal or corrected-to normal vision and reported no history of neurological disorders or dyslexia. The experiment was approved by the Cambridge Psychology Research Ethics Committee and volunteers were paid for their time and effort. (This experiment and its full details are described in Farahibozorg, 2018)

#### 2.1.2 Stimuli

The stimulus set included in our MEG analysis consisted of 250 uninflected words, including three categories of concrete words with strong visual, auditory and hand-action attributes (50 words per category), as well as two categories of emotional and neutral abstract words (50 words per category). For the purpose of this study, all the 250 words were pooled and a summary of their psycholinguistic variables is presented in Table 1. Concreteness ratings were obtained based on a word rating study (Farahibozorg, 2018) and CELEX Frequency, Orthographic Neighbourhood, Bigram and Trigram Frequencies were taken from the MCWord Database (Medler and Binder, 2005). Additional filler pseudowords were also included in the experiment, which are not assessed in this study.

**Table 1.** Psycholinguistic properties of stimuli included in EEG/MEG data analysis.

| | average ± standard deviation |
| --- | --- |
| **Number of Letters** | 5.68±1.56 |
| **CELEX Frequency** | 16.13±22.14 |
| **Orth Neighbourhood** | 3.78±4.81 |
| **Bigram Frequency** | 19008.54±9584.31 |
| **Trigram Frequency** | 1866.29±2278.84 |
| **Concreteness Rating** | 4.44±1.72 |

|  | **average ± standard deviation** |
| --- | --- |
| **Number of Letters** | 5.68±1.56 |
| **CELEX Frequency** | 16.13±22.14 |
| **Orth Neighbourhood** | 3.78±4.81 |
| **Bigram Frequency** | 19008.54±9584.31 |
| **Trigram Frequency** | 1866.29±2278.84 |
| **Concreteness Rating** | 4.44±1.72 |

#### 2.1.3 Procedure

The EEG/MEG experiment comprised four blocks presented in random order, and lasted approximately 90 minutes. We included 10-minute breaks between the blocks and short breaks every three minutes within each block. Each stimulus was presented for 150ms, with an average SOA of 2400ms (uniformly jittered between 2150 and 2650ms). Stimuli appeared as 30-point Arial font in black on a grey screen within a visual angle of 4 degrees in a slightly dimmed and acoustically shielded MEG chamber. One of the four blocks consisted of a lexical decision task and the remaining three blocks consisted of semantic target detection tasks. Half of the participants were randomly assigned to perform the lexical decision first and the other half performed semantic target detection blocks first. Details of these blocks were as follows:

1. Semantic target detection blocks: In each block, participants were presented with 250 words, as well as the filler items (overall 300 stimuli), in addition to 30 targets. They were asked to quietly read the strings of letters as they appeared on the screen and make button press responses with their left-hand middle finger when they saw a target on the screen. Each block had different targets which were selected from three groups of “non-citrus fruits”, “something edible with a distinctive odour” and “food that contains milk, flour or egg”. Block orders were randomised, and data acquired from the three blocks were pooled in the later EEG/MEG analyses in order to minimise possible question-specific effects.
2. Lexical decision task: Participants also performed a lexical decision task with the same 250 words, and 250 filler pseudowords to acquire response balance across stimuli (overall 500 stimuli). Participants were asked if “the following string of letters refers to a meaningful word” and they were asked to make button press responses with the index and ring fingers of their left hand for words and pseudowords, respectively. Only word stimuli were included in the subsequent EEG/MEG analyses.

### 2.2 Data Acquisition and Pre-processing

MEG and EEG data were acquired simultaneously using a Neuromag Vectorview system (Elekta AB, Stockholm, Sweden) and MEG-compatible EEG cap (EasyCap GmbH, Herrsching, Germany) at the MRC Cognition and Brain Sciences Unit, University of Cambridge, UK (Farahibozorg, 2018). MEG was recorded using a 306-channel system that comprised 204 planar gradiometers and 102 magnetometers. EEG was acquired using a 70-electrode system with an extended 10-10% electrode layout. EEG reference and ground electrodes were attached to the left side of the nose and the left cheek, respectively. ElectroOculoGram (EOG) was recorded by placing electrodes below and above the left eye (vertical EOG) and at the outer canthi (horizontal EOG). Electrocardiogram (ECG) was recorded by placing one electrode on the lower left rib and another electrode on the right wrist. Data were acquired with a sampling rate of 1000Hz and an online band-pass filter of 0.03 to 330Hz. During pre-acquisition preparations, positions of 5 Head Position Indicator (HPI) coils attached to the EEG cap, 3 anatomical landmark points (two ears and the nose) as well as approximately 50-100 additional points that covered most of the scalp were digitised using a 3Space Isotrak II System (Polhemus, Colchester, Vermont, USA) and later used for co-registration of EEG/MEG recordings with MRI data.

We applied signal space separation with its spatiotemporal extension implemented in the Neuromag Maxwell-Filter software to the raw MEG data in order to remove noise generated from sources distant to the sensor array (Taulu and Kajola, 2005). All remaining analyses were performed in the MNE-Python software package (Gramfort et al., 2014, 2013). Raw data were visually inspected for each participant, and bad EEG channels were marked and linearly interpolated. Data were then band-pass filtered using a finite-impulse-response (FIR) filter between 0.1 and 45 Hz. FastICA algorithm (Hyvarinen, 1999; Hyvärinen and Oja, 2000) was applied to the filtered data to remove eye movement and heartbeat artefacts. After ICA, data were divided into epochs from 300ms pre-stimulus to 600ms post-stimulus.

### 2.3 Source Estimation

We used L2-Minimum Norm Estimation (MNE) (Hämäläinen and Ilmoniemi, 1994; Hauk, 2004) for source reconstruction. Inverse operators were assembled based on a 3-layer Boundary Element Model (BEM) of the head geometry derived from structural MRI images, assuming sources perpendicular to the cortical surface (“fixed” orientation constraint). The MEG sensor configurations and MRI images were co-registered by matching the scalp digitisation points from the MEG preparation to the scalp surface reconstructed from individual MRI images. The noise covariance matrices for each individual and run were calculated for baseline intervals of 300ms. To do so, we used a list of methods from MNE python, ‘shrunk’, ‘diagonal_fixed’, ‘empirical’, ‘factor_analysis’, and the best estimator (‘shrunk’ in most cases) was selected using log-likelihood and cross-validation (Engemann and Gramfort, 2015). MNE-Python’s default SNR = 3.0 was used for evoked responses to regularise the inverse operator. The individuals’ results were then morphed to the standard average brain (fsaverage), yielding the time courses of activity for 20484 vertices for each subject and condition. It is noteworthy that the non-uniqueness of the EEG/MEG inverse problem leads to restricted spatial resolution, which may result in systematic mislocalisation of the genuine sources (Fuchs et al., 1999; Hauk et al., 2011; Molins et al., 2008), or more generally signal leakage between regions (Colclough et al., 2015; Palva et al., 2018; Wens et al., 2015; Williams et al., 2019).

### 2.4 Regions of Interest

Six regions of interest were defined using the anatomical masks provided from the Human Connectome Project (HCP) parcellation (Glasser et al., 2016), to represent the core semantic network as described in the introduction. As Figure 1a shows, this includes left and right ATLs (as defined in HCP: TGd, TGv, TE1a, anterior portions of TE2a and TE1m cut to terminate at the posterior extent of TE1a), left IFG (44, 45, 47l, p47r), left posterior temporal cortex (PTC, including posterior middle and inferior temporal gyri) (TE1p and posterior portions of STSvp, anterior inferior part of PH, and posterior portion of TE2p, all cut to terminate at the anterior limit of TE1p), left AG (PGi, PGp, PGs) and left primary visual area (PVA) (V1, V2, V3, V4).

**Figure 1.**
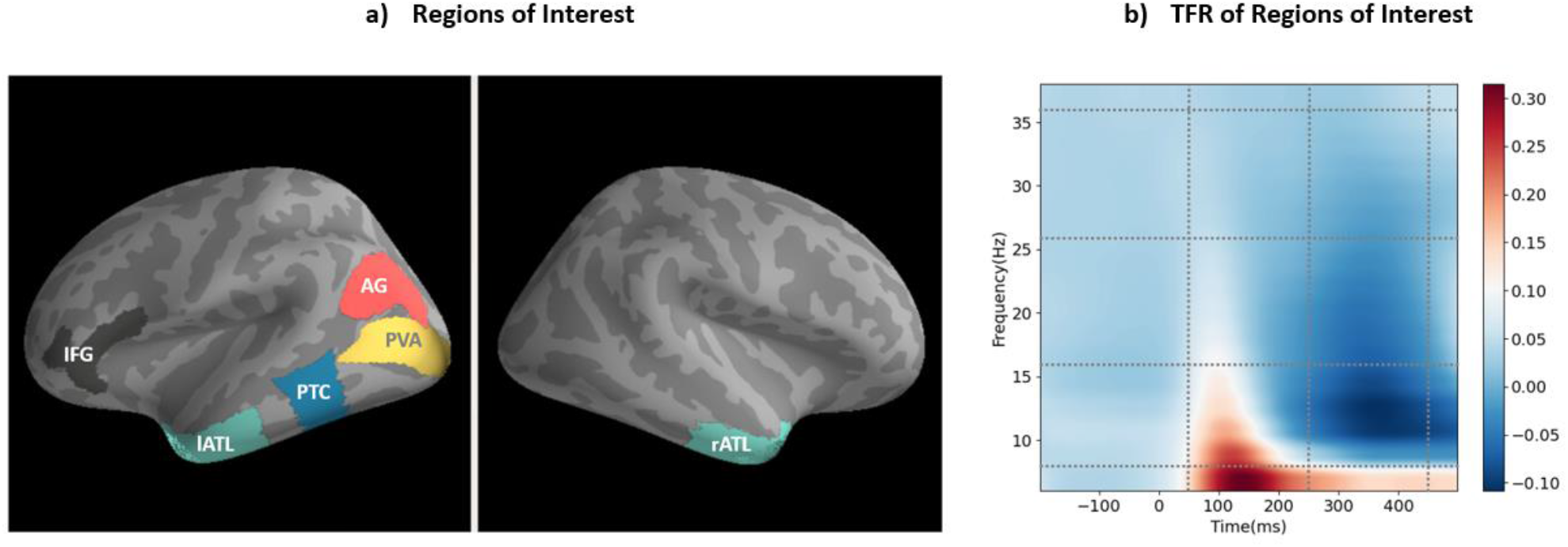
a) Regions of interest (ROIs) based on the semantic literature, b) Time-Frequency Representation (TFR) across all of ROIs, two tasks, and all participants.

### 2.5 Leakage

Source leakage is inherent in EEG/MEG source estimation due to the non-uniqueness of the inverse problem. Here, we provide a quantitative description of the source leakage among our ROIs. To have a better insight into the pattern of potential leakage, we computed the point spread and cross-talk functions (PSFs and CTFs; Hauk et al., 2011; Liu et al., 2002) of all the ROIs, to test how activity from one ROI leaks or spreads out to other regions. The general idea is to estimate the leakage from each ROI into all ROIs, relative to each ROI’s leakage into itself, in order to generate an ROI-to-ROI leakage matrix.

Thus, we defined the leakage index (LI) as follows:

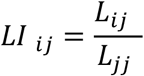

Where *L*_*ij*_ is leakage from *ROI*_*i*_ into *ROI*_*j*_ and *L*_*jj*_ is leakage from *ROI*_*j*_ into itself. Leakage can be described by PSFs, i.e., how each ROI leaks into the other ROIs, and CTFs, i.e., how all ROIs leak into one particular ROI. For the unweighted L2 minimum norm estimate, PSFs and CTFs are the same (its resolution matrix is symmetric, Hauk et al., 2019), and the leakage matrix, therefore, represents both types of leakage (similar to Farahibozorg et al., 2018).

Figure 2 presents PSFs and CTFs for our ROIs, as well as their associated leakage matrix. This shows that leakage varies across pairs of ROIs. In order to describe this variability, we will consider leakage indices between 0-0.2/0.2-0.4/0.4-0.6/0.6-0.8/0.8-1 as low/low-medium/medium/medium-high/high, which is reflected in the shading of the matrix cells. Leakage was medium and lower across all pairs of ROIs, and all leakage indices were below 0.5. Medium and high amounts of leakage can indicate that connectivity obtained from a pair of ROIs will be more affected by the limitations of the spatial resolution of the EEG/MEG source localisation and, thus, should be interpreted with more caution. Figure 2a confirmed that our ROIs produce most leakage in their vicinity. We will take individual leakage indices into account in our interpretation and discussion where appropriate. Please note that it is currently uncommon for non-methodological EEG/MEG studies to report this kind of information.

**Figure 2.**
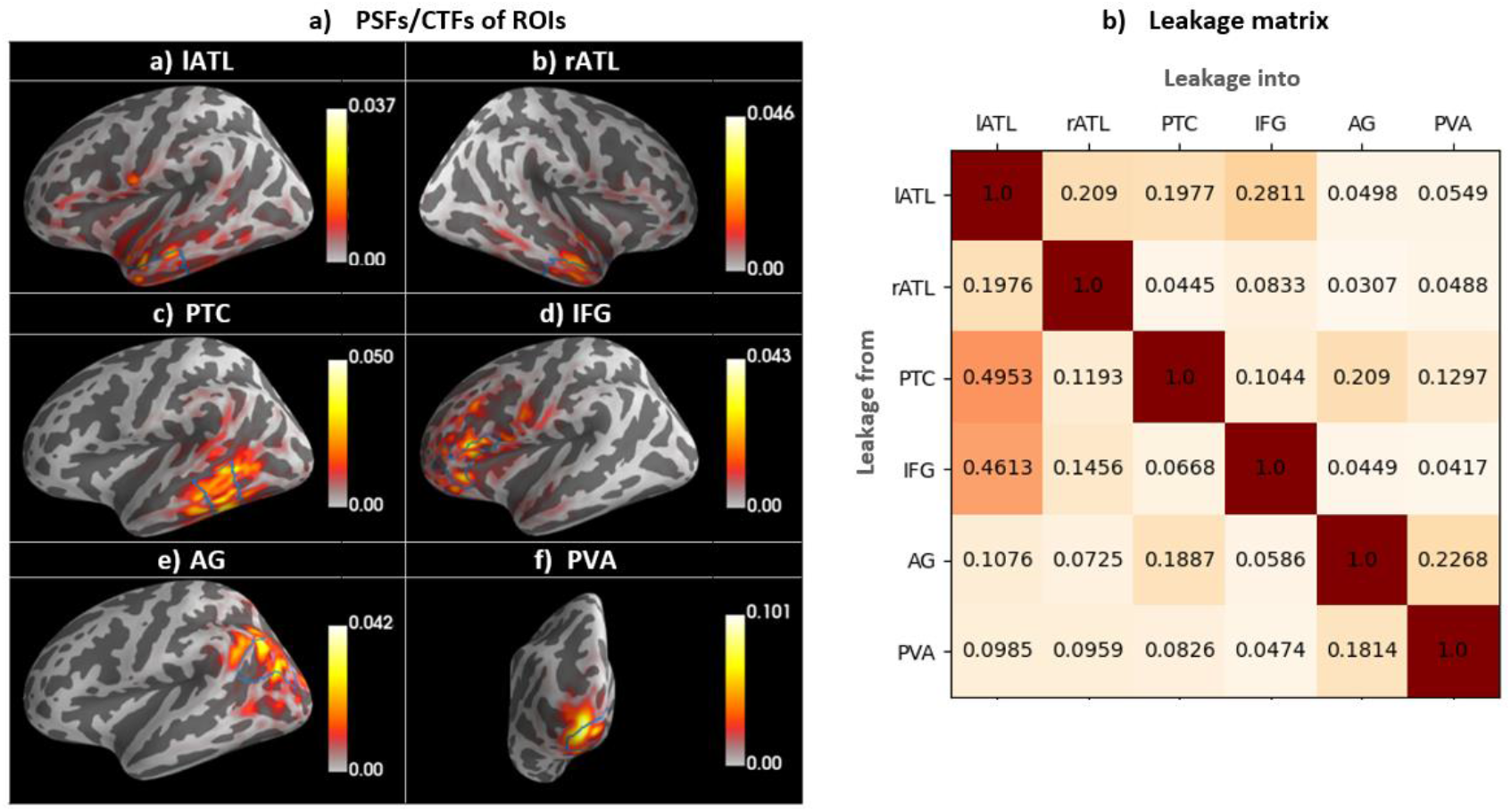
Leakage patterns. a) Grand average of PSFs/CTFs for each ROI, indicating how a real point source would leak to other regions (PSFs)/how all regions would leak to a particular ROI (CTFs). ROI borders are specified by solid blue lines. b) the leakage matrix; every column, corresponding to a single ROI, shows how much other regions leak into that ROI relative to what it leaks into itself.

### 2.6 Evoked Responses

The relevant trials for word stimuli were averaged in sensor space to obtain an evoked response per participant and task. Evoked responses were projected onto source space using L2-MNE (see above) and compared between the lexical and semantic decision tasks. For statistical analysis of the whole-cortex evoked responses, we used spatiotemporal cluster-based permutation tests (Maris and Oostenveld, 2007), accounting for multiple observations across vertices and time points. For this purpose, t-values were computed and thresholded with a t-value equivalent to p-value < 0.05 for a given number of observations, and randomisation was replicated 5000 times to obtain the largest random clusters. In addition to the whole-cortex analyses, activation time-courses were extracted from each ROI (using MNE Python’s “mean flip” option to account for varying source orientations within an ROI) and compared using cluster-based permutation tests per ROI.

### 2.7 Connectivity Analyses

Functional connectivity was estimated based on spectral coherence because it is sensitive to covariations of both phase and amplitude between two signals (Bastos and Schoffelen, 2016). We were also interested in potential zero-lag connectivity (e.g., between left and right ATLs), and therefore did not use the imaginary part of coherency or signal orthogonalisation (Colclough et al., 2015; Nolte et al., 2004). We will discuss any issues related to spatial resolution and leakage on the basis of our leakage analysis described above.

Whole-cortex seed-based connectivity was computed from each ROI. For this purpose, the ROI time-courses were extracted from each of the three blocks in the SD task, and from the LD task block. Magnitude-squared coherence was computed between each ROI time course and every vertex in the brain, for four different frequency bands and two time windows. The results were averaged across the three SD blocks for comparison with LD. This helped ensure that our coherence estimation is not biased due to different numbers of trials between the SD and LD task (Bastos and Schoffelen, 2016). In order to choose frequency bands and time intervals of interest in an unbiased manner, we present the time-frequency representation of our dataset across all conditions, participants, and ROIs in Figure 1b. Based on prominent features of this time-frequency representation, i.e., peaks, increases and decreases of activity, we selected an early (50-250ms) and late (250-500ms) time window and computed coherence in four frequency ranges, namely theta (4-8Hz), alpha (8-16Hz), beta (16-26Hz), and gamma (26-36Hz).

## 3 Results

### 3.1 EEG/MEG Behavioural Results

For the lexical decision task, the average and standard deviation reaction times were 0.66±0.069s and response accuracies were 95.09±3.96%. For the semantic target detection blocks, the target detection accuracy and reaction times were 0.90±0.11 and 0.99±0.22s, respectively.

### 3.2 Whole-Cortex Evoked Analysis

Most previous investigations into the neuronal basis of semantics using EEG/MEG and source estimation based their main conclusion on ROI-based analysis approaches. While this increases statistical sensitivity, it raises questions with respect to the spatial specificity of the reported effects, especially since the limited spatial resolution and possible mis-localisation of EEG/MEG source estimation are well-documented (Hauk et al., 2019; Molins et al., 2008). However, whole-cortex analyses in different latency and frequency ranges can be hard to present and interpret. In the following, we will present a hybrid approach that starts with whole-cortex results followed by ROI-based results. Our main conclusions will be based on the commonalities of the two analyses, and we will discuss any discrepancies, where appropriate.

To track task modulation of brain activation over time, we first compared evoked brain activity between our two tasks using whole-cortex cluster-based permutation tests in five non-overlapping time windows of 100ms duration starting at 50ms after stimulus onset. These brain dynamics were then analysed in more detail using an ROI analysis. The results of the whole-cortex evoked analysis are displayed in Figure 3. The colour-coding indicates the duration of significant activation within each time window. Importantly, task differences were already apparent in the first time window (50-150ms) and remained significant throughout the first three windows, until 350ms.

**Figure 3.**
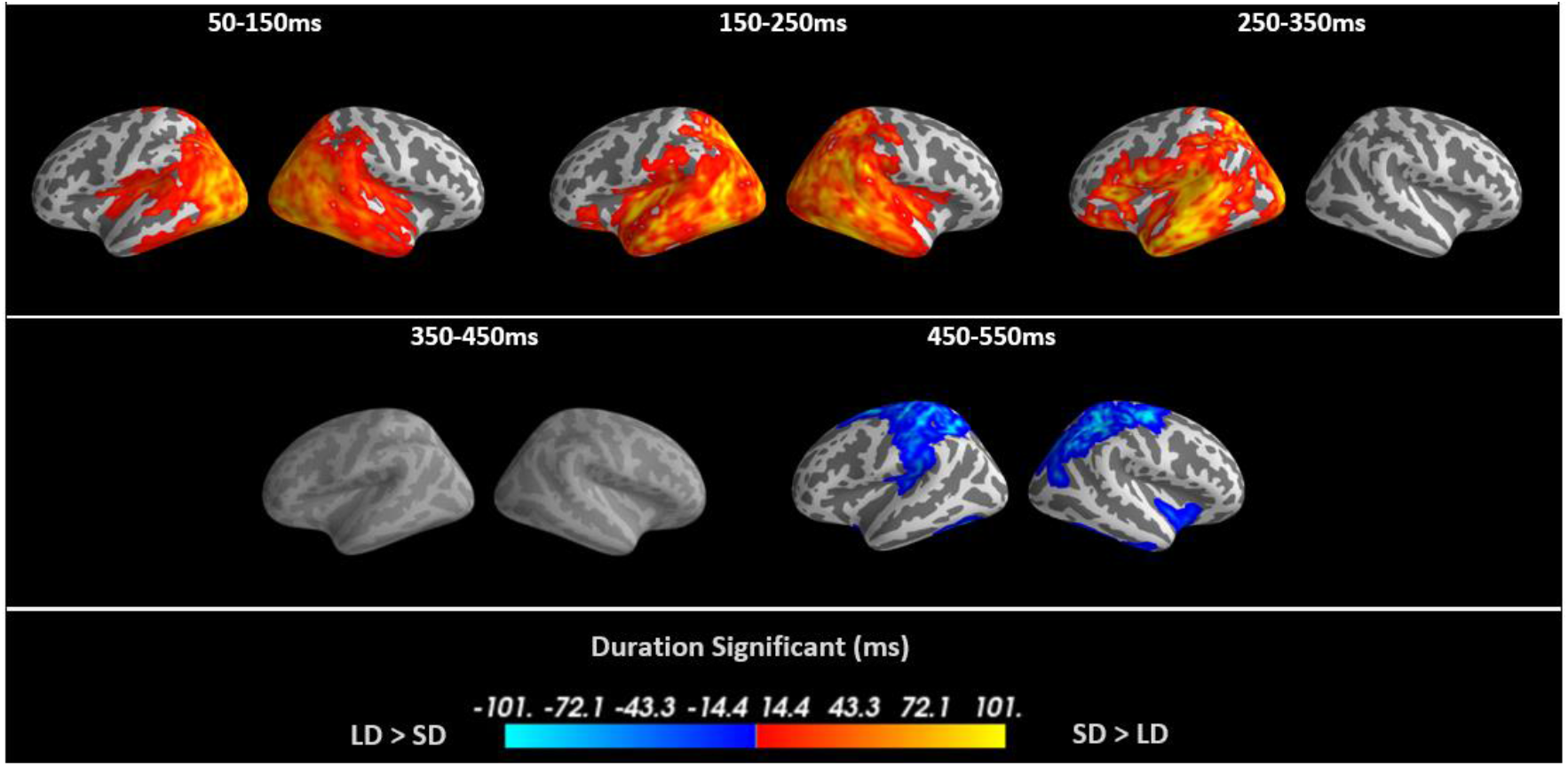
Spatiotemporal cluster-based permutation test contrasting the evoked responses of Semantic Decision (SD) and Lexical Decision (LD) in five time windows. The first three time windows showed significantly greater activation for SD than LD, across the semantic network, with early effects in occipital and temporal cortex and later differences in frontal cortex. These changes are initially bilateral and later left-lateralised. The last time window demonstrates significant activation for LD in motor regions, likely caused by the need for more frequent responses.

The semantic decision task produced higher levels of activation compared to the lexical decision task up to 350ms. The earliest task differences were predominantly in bilateral posterior brain areas, but differences were already present in inferior parietal and anterior temporal brain regions. Between 150-250ms, task modulations spread further into anterior temporal and parietal regions in both hemispheres. After 250ms, activation was strongly left-lateralised and included left inferior frontal regions. There were no significant task differences between 350-450ms. The lexical decision task produced larger activation than the semantic decision task between 450-550ms, presumably due to the button-press-related finger movements for words in the lexical decision task.

### 3.3 ROI Activation Time-courses

Figure 4 presents the millisecond-by-millisecond time courses of evoked brain activity for our selection of ROIs. Averaged time courses across participants are shown for each individual task and their subtraction, alongside the t-values of their statistical comparison. Shaded areas highlight the latency ranges with significant task differences using cluster-based permutation tests. The earliest task differences occurred in PVA and AG from 60 and 65ms. Note that the leakage indices for these two regions (Figure 2b) were about 0.2, and their time courses are similar (Figure 4 e and f). Therefore, we cannot rule out the possibility that these results reflect leakage effects, i.e., are due to the same neuronal sources in posterior brain areas. These early effects were followed by differences in PTC and lATL at 186 and 189ms. We also found marginally significant task differences at later latencies in PVA at 300ms, and IFG and PTC at 309ms. We observed no statistically significant task difference in rATL at any latency.

**Figure 4.**
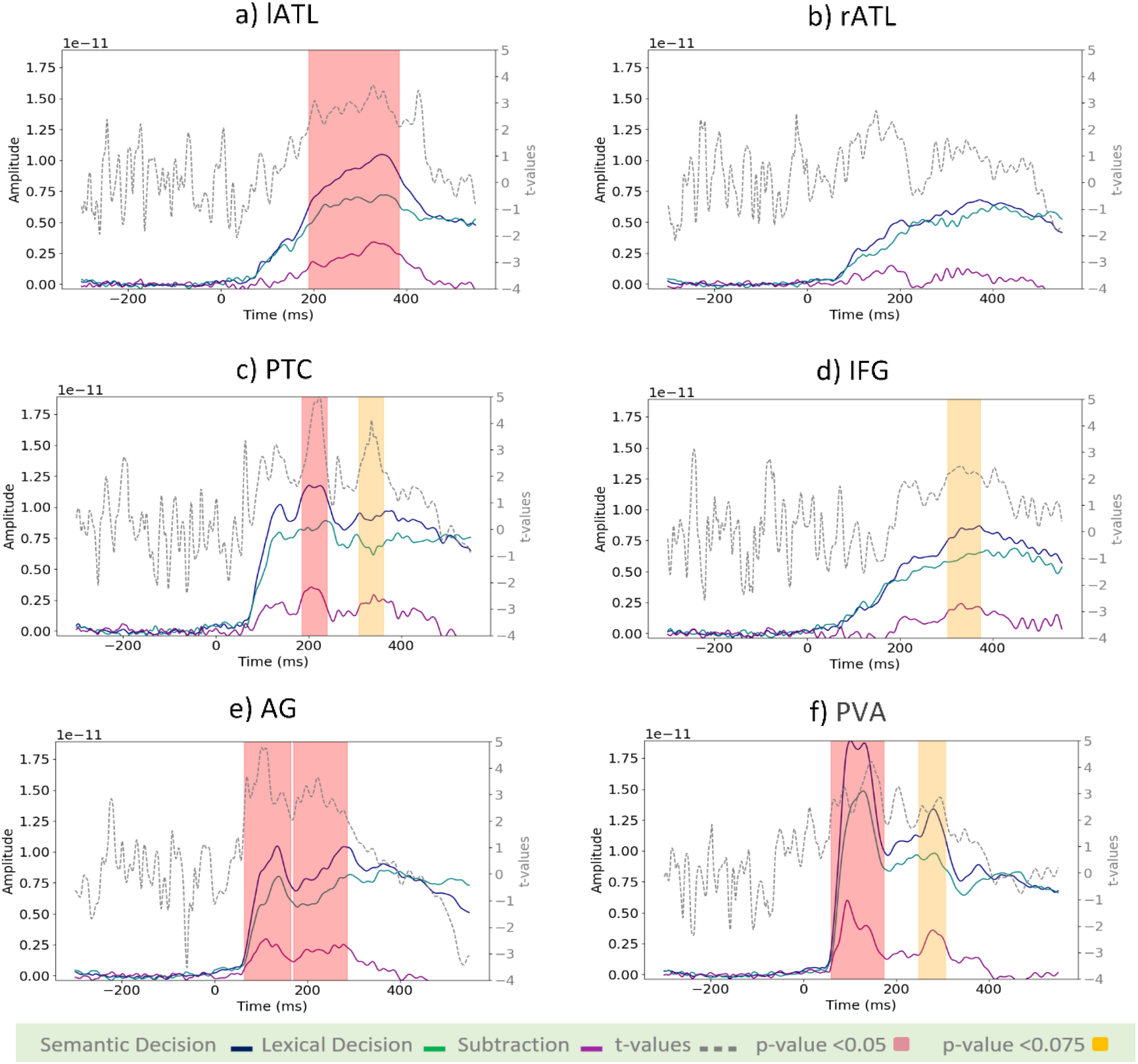
Activation time-course of ROIs for SD, LD, SD-LD, and t-values. The left-hand side axis represents source amplitudes, the right-hand side axis shows t-values for the comparison of SD and LD, and the horizontal axis represents time in milliseconds. t-values corresponding to p-value <0.05 have been highlighted in red, and those with p-value <0.075 in yellow.

### 3.4 ATL Laterality

We explicitly tested the laterality of ATL involvement. Figure 5a shows the main effects of task, laterality, and their interaction using a two-way repeated-measures ANOVA. To understand this interaction, six planned comparisons were run. Figure 5b shows separate activation time courses for left and right ATL for lexical and semantic decision tasks, respectively. Figure 5c displays the contrasts that yielded significant results. Figure 5d presents a summary statistical analysis of activation averaged in the time window 150-400ms. This analysis demonstrated that the task effects in the ATLs were driven by larger activation in the left, but not right, ATL for the semantic decision task than the lexical decision task ([SD[lATL]-SD[rATL]: (t=4.13, p< 0.001)], [SD[lATL]-LD[lATL]: (t=3.00, p< 0.01)], [SD[lATL]-LD[rATL]: (t=4.76, p< 0.001)], [SD[rATL]-LD[lATL]: (t=-0.66, p> 0.50)], [SD[rATL]-LD[rATL]: (t=1.24, p> 0.20)], [LD[lATL]-LD[rATL]: (t=1.82, p> 0.08)]). Thus, the left and right ATL responded similarly to the less demanding lexical decision task, yet the increased requirements of the semantic decision task were met by a greater response from the left ATL, in particular.

**Figure 5.**
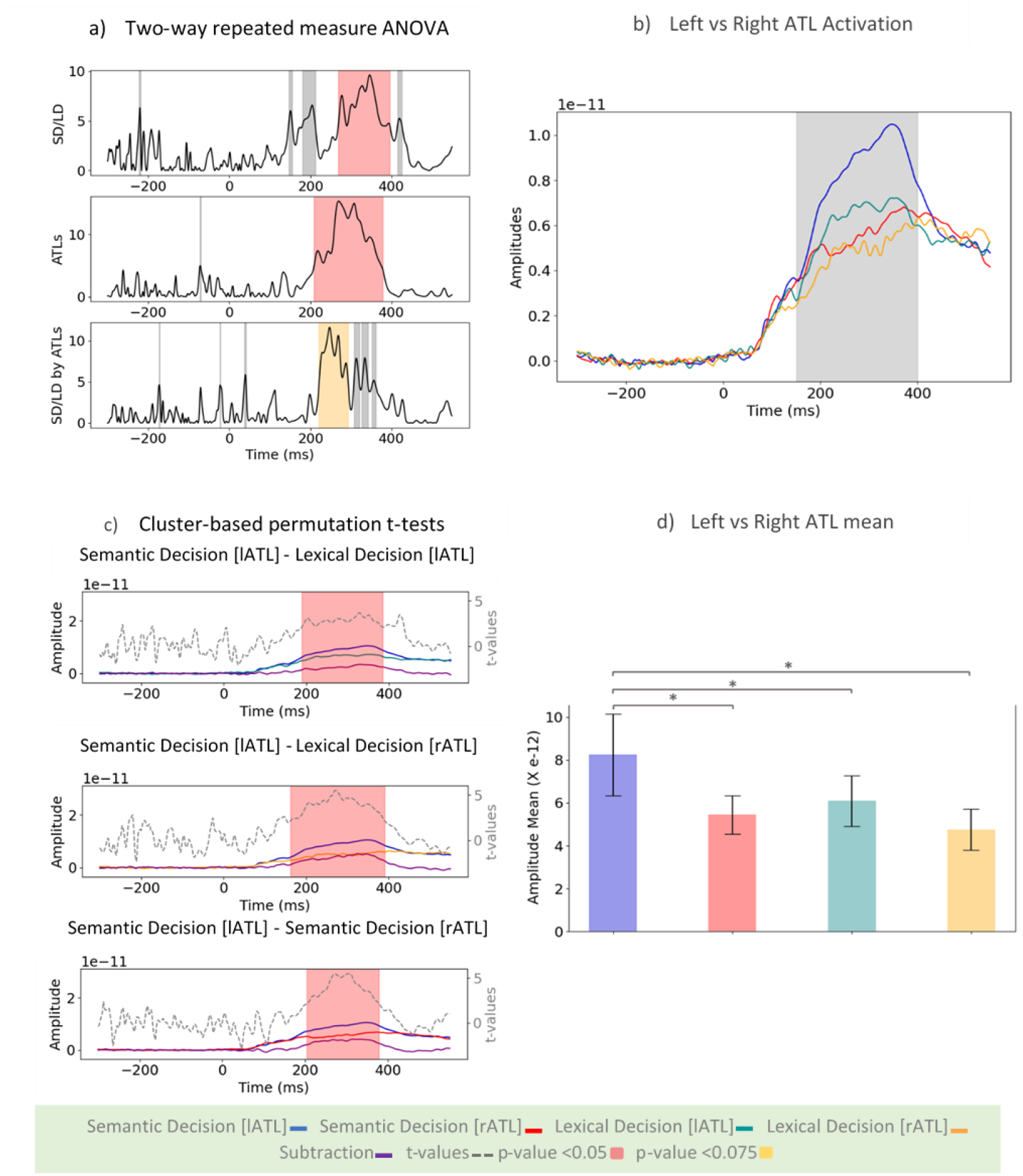
Laterality of task modulation in ATLs. a) the effect of Task (SD vs. LD), Laterality (left vs. right), and their interaction using a two-way repeated-measures ANOVA. b) activations of the left and right ATLs in SD and LD. c) all comparisons between the left and right ATL activation in each task that reach significance using cluster-based permutation test (three out of six). t-values corresponding to p-value <0.05 have been highlighted in red, and those with p-value <0.075 in yellow. d) average activation of the left and right ATLs in SD and LD tasks in the time range of 150 to 400ms (shaded grey area in panel b), chosen based on the interaction results.

### 3.5 Connectivity Analysis: Whole-Cortex Seed-Based Connectivity

We studied task modulation of functional connectivity in the semantic network with a whole-cortex seed-based analysis, followed by ROI analyses. The seed-based analysis determined the coherence between our ROIs and all other vertices in the brain in two time windows (early 50-250ms and late 250-450ms) and four frequency bands (theta, alpha, beta, gamma). The whole-cortex seed-based connectivity results are presented in Figure 6. Statistical significance was assessed based on whole-cortex cluster-based permutation tests. We found no significant effects in the theta band, which may be too slow to reflect the short-lived processes involved in semantic single-word processing.

**Figure 6.**
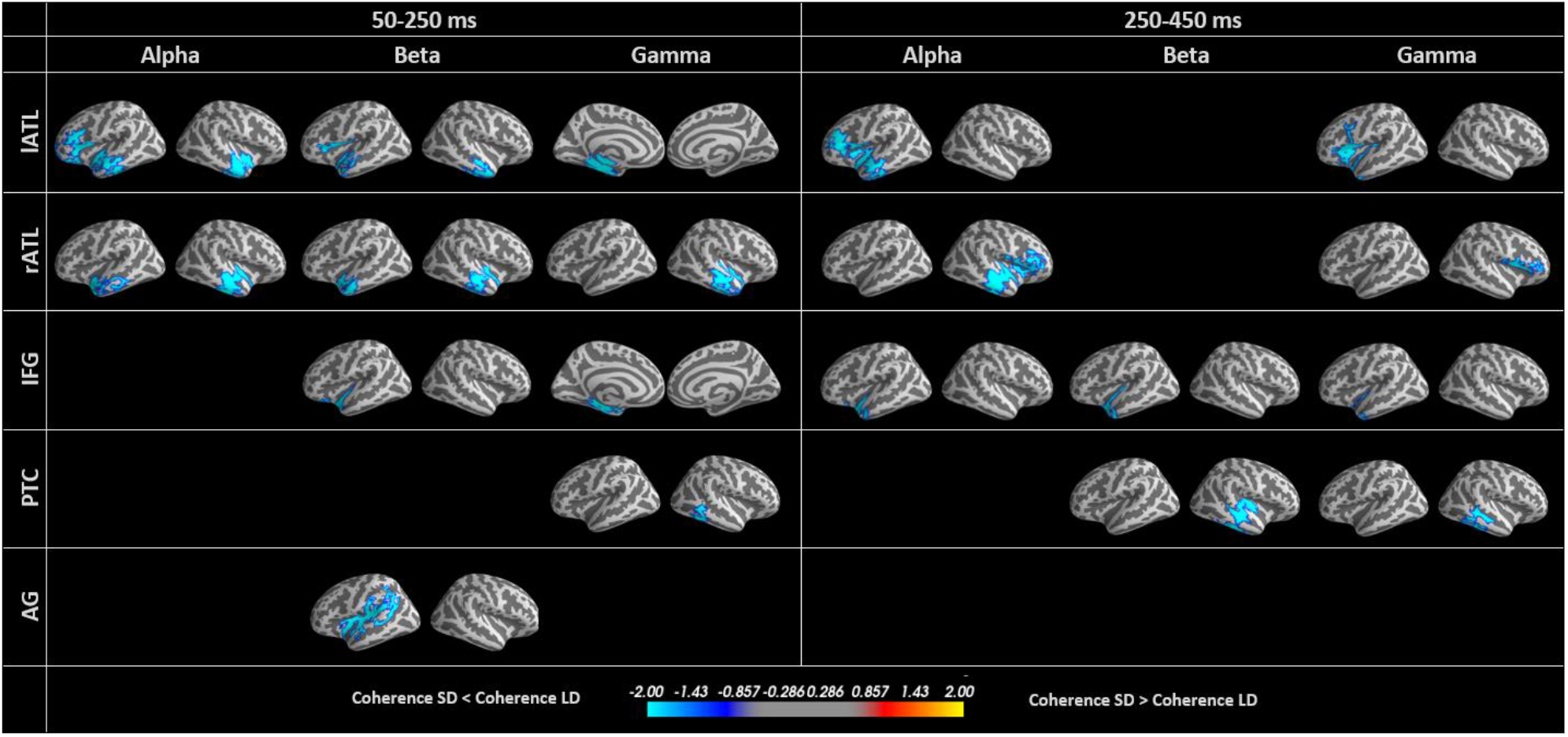
Whole-cortex seed-based connectivity differences between the semantic and lexical decision tasks.Connectivity between lATL and rATL was modulated in the first time window, and connectivity between lATL and left IFG was modulated across both time windows. PTC demonstrated changes in connectivity to homologous regions, and AG showed connectivity changes outside the semantic network.

The ROI labels in the left column indicate the seed region. Note that each seed region strongly “leaks” into itself (Figure 2b), and therefore we can expect high coherence values within each seed region for the individual tasks. However, if these values are similar for lexical and semantic decisions, they will not produce significant effects in the subtraction (or statistical comparison). Significant effects in seed regions may still occur due to other factors (e.g., noise levels), but we will not interpret them in terms of functional connectivity. We will take possible leakage into account in the interpretation of functional connectivity (see Figure 2).

Interestingly, our functional connectivity analyses generally revealed larger coherence values for lexical compared to semantic decisions, which may appear counterintuitive or in contrast to our evoked analyses showing larger activation for semantic decisions. However, this could be explained by larger trial-by-trial variability and higher desynchronisation leading to lower coherence in the more demanding task. We will come back to this issue in the discussion.

In the early time window, we observed significant task differences in functional connectivity between the left and right ATL in alpha and beta bands, as well as from AG along the Sylvian fissure, including anterior superior temporal lobe. The gamma band demonstrated significant modulation of connectivity between the IFG seed and left ATL, and between PTC and an approximately homologous area in the right hemisphere.

In the late time window, we found significant task modulation of connectivity between lATL and left IFG, as well as rATL and right IFG, but not between lATL and rATL as found in the early time window.

AG did not demonstrate any connectivity differences in this time window, while PTC showed differential connectivity with an area of right middle temporal lobe in the beta and gamma bands. In the gamma band, there was task modulation of the connectivity between the lATL and IFG.

Thus, most differences in the connectivity of the lexical and semantic decision tasks involved the ATLs, with task modulation principally affecting the connectivity between left and right ATL at early stages, and later, between IFG and ATL.

### 3.6 Connectivity Analysis: Between-ROIs Connectivity

As with the evoked analysis, we sought to corroborate our whole-cortex seed-based analysis using an ROI approach. The results displayed in Figure 7 confirm significant task-dependent connectivity between lATL and rATL for alpha and beta bands in the early latency window, as well as between lATL and IFG for alpha and beta bands in the late window. The gamma band also showed task modulation of connectivity between lATL and rATL and between rATL and PTC in the early window, and between lATL and IFG in the late window. Furthermore, the gamma band produced significant connectivity differences between AG and IFG, which is the only case where coherence values are larger in the semantic compared to the lexical decision task. In the late window, the gamma band also showed a connection between AG and PTC.

**Figure 7.**
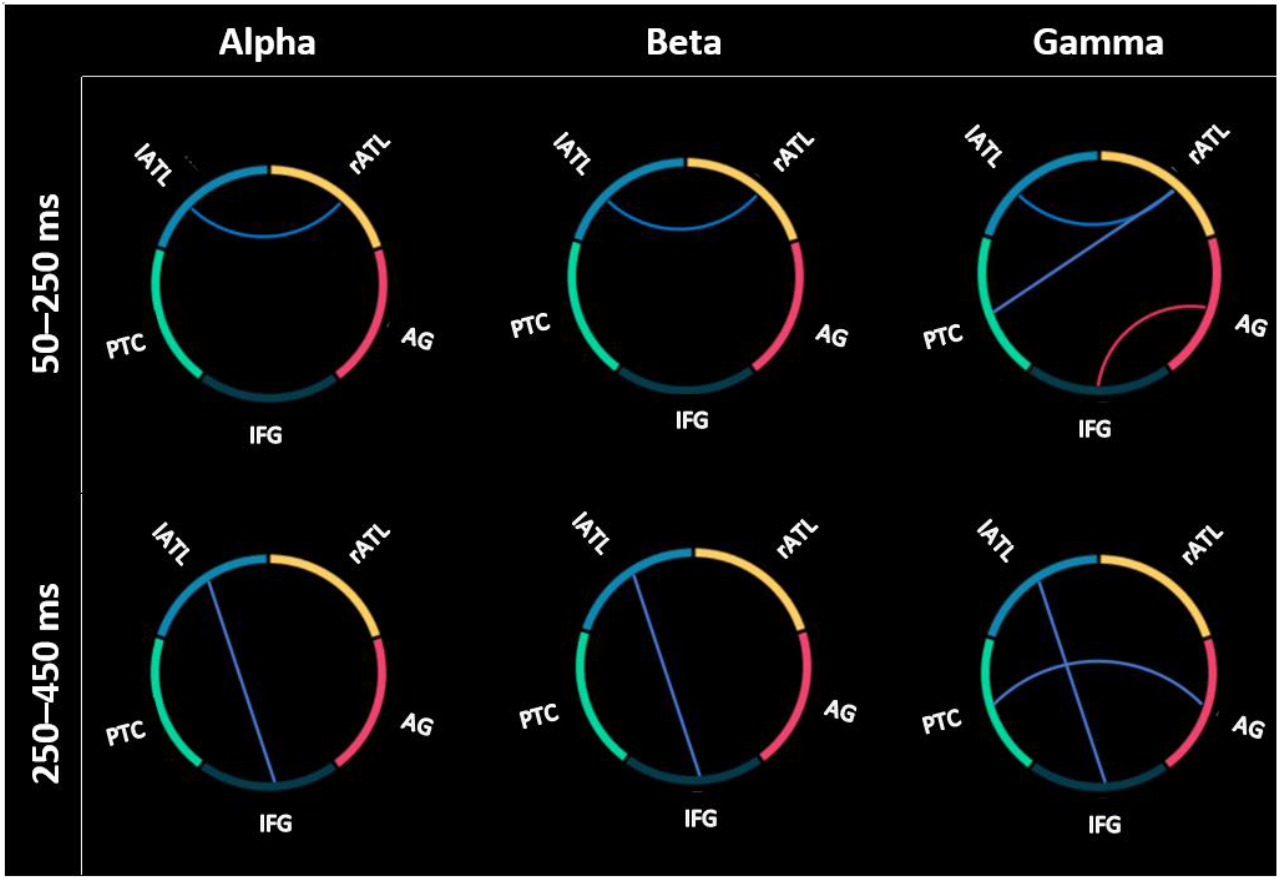
Significant differences in connectivity in the semantic and lexical decision tasks between the semantic ROIs. In the early time window, connectivity between left and right ATL was modulated, and in the later time window the connectivity between lATL and IFG was modulated.

This analysis confirmed the early task differences in the connectivity between left and right ATL and the later differences in their connectivity with IFG, found in the whole-cortex seed-based analysis. Pushing the semantic system modulates the connectivity between core regions of the semantic network; the left and right ATLs and the IFG.

## 4 Discussion

Semantic cognition critically depends upon interaction across a distributed network, yet very few studies have elucidated the connectivity of this semantic network using high temporal resolution techniques. Here we investigated the effects of increasing semantic task demands on spatiotemporal brain activity and functional connectivity in the semantic network estimated from combined EEG/MEG data. We asked how the need for greater semantic cognition modulates responses in the semantic brain network over time, how it affects connectivity among putative semantic representation and control regions, and specifically how it modulates the laterality and connectivity of left and right ATLs. In our whole-cortex evoked analysis, we observed task differences in bilateral posterior brain regions already present within the 50-150ms time window, with greater semantic demand resulting in greater activation across much of the semantic network until 350ms. Early task differences involved inferior parietal and temporal brain regions, which spread further into anterior temporal and parietal regions in both hemispheres between 150-250ms. After 250ms, the task-modulated evoked activation became left-lateralised and also spread to left inferior frontal regions. Laterality effects in ATL were driven by larger activation in left ATL in the more semantically demanding task. Regardless of the precise assessment method used (whole-cortex seed-based or ROI-based functional connectivity analyses), functional connectivity was modulated by task in multiple frequency bands and time windows, especially between left and right ATL at early latencies in the alpha and beta bands, and between left and right ATL and IFG in later time windows in the alpha and gamma bands. These effects reflected larger desynchronisation in SD compared to LD. Our results indicate that semantic representation and control processes dynamically interact within the first few hundred milliseconds of written word processing, and confirm that the ATL has a central role in the semantic brain network.

Spatiotemporal evidence for the interplay of representation and control processes in dynamic semantic brain networks is still scarce. Here, we contrasted a more semantically-demanding (semantic decision) task with a less semantically-demanding (lexical decision) task on the same set of well-matched word stimuli. This task contrast does not allow us to unambiguously disentangle representation and control processes, but provides critical information as to the dynamics of the semantic network overall, including the interaction between putative semantic representation and control regions. In addition, the temporal and spectral information presented provides novel insights that will be the basis for future studies on this issue. Importantly, our study included some methodological advancements with respect to the majority of previous EEG/MEG studies on semantic word processing. First, we used combined EEG and MEG recordings in order to optimise spatial resolution for source estimation using individual realistic head modelling (Hauk et al., 2019; Molins et al., 2008). Second, we present both conventional evoked responses, as well as functional connectivity results in source space in the same study. Third, we provide both whole-cortex and ROI-based results to strike a trade-off between sensitivity and spatial specificity. Fourth, we explicitly evaluated the spatial resolution (“leakage”) of our ROIs as a basis for a critical interpretation of our source estimation results (Hauk et al., 2019). We hope that it will become standard in the EEG/MEG literature to report the relevant leakage indices (or similarly informative measures) in the future.

Whilst our evoked analysis showed the expected pattern of more activation for the more semantically-demanding task, the opposite seemed to be the case in our functional connectivity analyses, where LD showed larger coherence values compared to SD. However, coherence values reflect the variability of amplitude and phase across trials (Bastos and Schoffelen, 2016; Lachaux et al., 1999). This LD task is less demanding, resulting in lower reaction times and presumably lower variability across trials, therefore possibly resulting in larger coherence values. Furthermore, Hanslmayr et al. (2012) linked neural desynchronisation to information theory, suggesting that more processing demands can result in larger desynchronization within neuronal populations, in our case resulting in lower coherence values in the SD task, which would fit our observation of lower coherence values in the SD task. We, therefore, simply conclude that our coherence effects reflect task modulation of functional connectivity in the semantic brain network, without further interpretation of the direction of the difference. Note that the use of a spectral connectivity method does not immediately imply that our observed effects reflect specific mechanisms, such as neuronal “oscillations”. Indeed, the fact that some effects, such as connectivity between left and right ATLs, occurred in multiple frequency ranges suggests that the underlying neuronal processes may be a broadband phenomenon. The more specific neurophysiological mechanism will have to be studied in more detail in the future.

Large task modulations were found throughout the putative semantic network; in bilateral ATLs, IFG, PTC and visual cortices. Engagement of the semantic network is not all or nothing; the information accessed and employed depends upon task demands, even in early word processing (Chen et al., 2015, 2013; Jackson, 2021; Jefferies, 2013; Strijkers et al., 2015). We found early task modulation in visual and inferior parietal areas followed by temporal lobe structures, in particular left anterior temporal lobe and posterior temporal cortex, and then inferior frontal regions. These regions are critical for demanding semantic cognition and are recruited flexibly based on task demands. In particular, the connectivity between left and right ATL and between ATL and IFG supports demanding semantic cognition. This is highly compatible with prior functional connectivity assessments of the semantic network, including the compensatory effects of connectivity between left and right ATLs after transcranial magnetic stimulation (Chiou et al., 2018; Chiou and Lambon Ralph, 2019; Farahibozorg et al., 2019; Jackson et al., 2016; Jung and Ralph, 2019). The selection of task-relevant, and inhibition of task-irrelevant, semantic information is hypothesised to require the interaction of control regions (which represent the task context, including IFG) and representation areas (where task-independent semantic representations are stored, hypothesised to rely principally on the ATLs, Jackson, 2021; Jefferies, 2013; Lambon Ralph et al., 2016). Thus, the differential connectivity between the ATLs and the IFG in the semantically demanding task may reflect the additional interaction required to access the specific subset of features required to answer the difficult semantic decisions.

Very early changes were identified in visual and parietal regions. As described before, the spatial resolution of EEG/MEG does not allow an interpretation at the same level of spatial detail as for fMRI, e.g., with respect to the exact Brodmann areas. This was confirmed by our leakage analysis. However, the high temporal resolution of EEG/MEG allows us to conclude that task modulations that occurred in “early visual areas” (as they are sometimes called in the fMRI literature without timing evidence (e.g. Basti et al., 2019; Mur et al., 2012)) indeed reflect early brain processing, rather than recurrent activation flow (e.g. Lamme and Roelfsema, 2000).

Early task modulation effects were also identified in the AG, a region with a debated role in semantic cognition (Binder et al., 2009; Humphreys et al., 2015; Noonan et al., 2013). Whilst our leakage analysis suggests that AG effects are unlikely due to leakage from PVA, the point-spread and cross-talk functions in Figure 2 indicate that there could be significant leakage from higher level visual areas posterior to AG but anterior to PVA. Nevertheless, some previous MEG studies have reported AG involvement in semantic processes (e.g., Lewis et al., 2015; Williams et al., 2017). However, in our connectivity analysis, AG does not show rich connectivity with other semantic areas, especially not in the temporal lobes. Additionally, these effects are very early, in parallel with visual areas and prior to any other semantic region. Our ROI-based connectivity analysis (Figure 7) revealed connectivity modulation between AG and IFG in the early time window, although this was in the opposite direction to all other connectivity differences. Some previous neuroimaging studies have suggested that AG may serve semantic representation (Binder et al., 2009) or control functions (Noonan et al., 2013, but see Jackson 2021), although these assessments are plagued by questions of how to interpret differences in the context of difficulty-dependent deactivation in this region (Humphreys et al., 2015; Humphreys and Lambon Ralph, 2014). Indeed, the AG is consistently found as part of the default mode network (Buckner and DiNicola, 2019) and may play a role in attentional processes. For instance, our early task effects in AG could reflect a change from readiness during rest to the engagement of task networks by this area, resulting in an early increase or “boost” of attentional resources towards the visual word form or the semantic network. Alternatively, a similar function could be achieved by task-positive inferior parietal regions (Duncan, 2010) and the current effects misattributed to the AG region. This hypothesis can be tested in future studies using more fine-grained experimental paradigms.

Our evoked analysis revealed task modulation in ATL starting prior to 200ms. Previous EEG, MEG and behavioural studies have suggested that semantic information becomes available in visual word processing around this latency (Amsel et al., 2013; Hauk et al., 2012; Pulvermüller et al., 2009), and some EEG/MEG studies have reported activity in ATL regions (Bemis and Pylkkänen, 2013; Dhond et al., 2007; Farahibozorg et al., 2019; Marinkovic et al., 2014; Mollo et al., 2017; Westerlund and Pylkkänen, 2014). This task effect was clearly left-lateralised in our evoked data, which is consistent with findings from neuropsychological and neuroimaging literature that left ATL shows a preference for linguistic stimuli and tasks (Rice et al., 2015a, 2015b). However, connectivity between the ATLs was significant in this early time window, highlighting the possibility of a critical role for the right ATL. Our results also indicate that areas that do not show a significant activity effect can still be part of a distributed network. Indeed, the laterality of the evoked responses changed over time suggesting an interpretation of the necessity of a single ATL may be an oversimplification of a dynamic, recurrent system. This significant functional connectivity in three frequency bands (alpha, beta and gamma) demonstrated that evoked and spectral responses carry independent information.

The connectivity between ATLs and the IFG also varied across time, with significant effects in the later time window. The PTC, another putative semantic control region, was engaged both at a similar time to the IFG (around 300ms) and at an earlier time point (around 200ms, with the ATL response). The relative timings of the putative semantic control and representation regions are informative as to their possible interactions. To date, it has been hard to separate the role of IFG and PTC (Jackson, 2021; Jefferies, 2013; Lambon Ralph et al., 2016) and their differential timings could be informative; e.g. could PTC be involved earlier? However, we cannot rule out the possibility that we cannot distinguish the control-related PTC changes from nearby regions engaged in semantic representation, due to the nature of the task manipulation. Perhaps the responses at the two different time points reflect these different elements of semantic cognition, with an early sweep through PTC before the semantic control regions are active. Indeed, although PTC demonstrated task modulated evoked responses, no clear changes in connectivity were identified. This could be a result of the particular connectivity measure chosen, or due to high levels of connectivity with other semantic regions across both tasks.

In conclusion, our results suggest that semantic task demands modulate visual word processing before 100ms in posterior visual (and perhaps attentional) areas, followed by modulation of multimodal semantic regions; first ATLs and PTC, and then IFG, allowing the context-appropriate extraction of task-relevant semantic features critical for response selection. Our conclusions required the high temporal and reasonable spatial resolution of combined EEG and MEG measurements, as well as the combination of evoked and functional connectivity analyses. Our results raised several questions about the precise mechanisms of the interaction of semantic control and representation, and provide a valuable base to address them in future EEG/MEG studies. In particular, the spatiotemporal resolution of combined EEG/MEG recordings together with sophisticated multivariate and multi-dimensional connectivity methods will be required to characterise dynamic semantic brain networks in more detail (Anzellotti and Coutanche, 2018; Basti et al., 2020, 2019; Kietzmann et al., 2019).

## 5 Acknowledgements

This work was supported by intramural funding from the Medical Research Council UK (MC_UU_00005/18), a British Academy Postdoctoral Fellowship awarded to R.L.J. (no. pf170068), and Cambridge University international scholarships awarded to S.R. and S.R.F. S.R.F. is supported by Wellcome Trust (215573/Z/19/Z, 203139/Z/16/Z). For the purpose of open access, the author has applied a CC BY public copyright license to any Author Accepted Manuscript version arising from this submission.

## References

Acosta-Cabronero, J., Patterson, K., Fryer, T.D., Hodges, J.R., Pengas, G., Williams, G.B., Nestor, P.J., 2011. Atrophy, hypometabolism and white matter abnormalities in semantic dementia tell a coherent story. Brain 134, 2025–2035.

Alam, T.R.J.G., Krieger-Redwood, K., Evans, M., Rice, G.E., Smallwood, J., Jefferies, E., 2021. Intrinsic connectivity of anterior temporal lobe relates to individual differences in semantic retrieval for landmarks. Cortex 134, 76–91.

Amsel, B.D., Urbach, T.P., Kutas, M., 2013. Alive and grasping: Stable and rapid semantic access to an object category but not object graspability. Neuroimage 77, 1–13.

Anzellotti, S., Coutanche, M.N., 2018. Beyond functional connectivity: investigating networks of multivariate representations. Trends Cogn. Sci. 22, 258–269.

Badre, D., Poldrack, R.A., Paré-Blagoev, E.J., Insler, R.Z., Wagner, A.D., 2005. Dissociable controlled retrieval and generalized selection mechanisms in ventrolateral prefrontal cortex. Neuron 47, 907–918.

Basti, A., Mur, M., Kriegeskorte, N., Pizzella, V., Marzetti, L., Hauk, O., 2019. Analysing linear multivariate pattern transformations in neuroimaging data. PLoS One 14.

Basti, A., Nili, H., Hauk, O., Marzetti, L., Henson, R.N., 2020. Multi-dimensional connectivity: a conceptual and mathematical review. Neuroimage 117179.

Bastos, A.M., Schoffelen, J.-M., 2016. A tutorial review of functional connectivity analysis methods and their interpretational pitfalls. Front. Syst. Neurosci. 9, 175.

Bemis, D.K., Pylkkänen, L., 2013. Basic linguistic composition recruits the left anterior temporal lobe and left angular gyrus during both listening and reading. Cereb. Cortex 23, 1859–1873.

Binder, J.R., Conant, L.L., Humphries, C.J., Fernandino, L., Simons, S.B., Aguilar, M., Desai, R.H., 2016. Toward a brain-based componential semantic representation. Cogn. Neuropsychol. 33, 130–174.

Binder, J.R., Desai, R.H., Graves, W.W., Conant, L.L., 2009. Where is the semantic system? A critical review and meta-analysis of 120 functional neuroimaging studies. Cereb. Cortex 19, 2767–2796.

Buckner, R.L., DiNicola, L.M., 2019. The brain’s default network: updated anatomy, physiology and evolving insights. Nat. Rev. Neurosci. 20, 593–608.

Chen, Y., Davis, M.H., Pulvermüller, F., Hauk, O., 2015. Early visual word processing is flexible: Evidence from spatiotemporal brain dynamics. J. Cogn. Neurosci. 27, 1738–1751.

Chen, Y., Davis, M.H., Pulvermüller, F., Hauk, O., 2013. Task modulation of brain responses in visual word recognition as studied using EEG/MEG and fMRI. Front. Hum. Neurosci. 7, 376.

Chiou, R., Humphreys, G.F., Jung, J., Lambon Ralph, M.A., 2018. Controlled semantic cognition relies upon dynamic and flexible interactions between the executive ‘semantic control’and hub-and-spoke ‘semantic representation’systems. cortex 103, 100–116.

Chiou, R., Lambon Ralph, M.A., 2019. Unveiling the dynamic interplay between the hub-and spoke-components of the brain’s semantic system and its impact on human behaviour. Neuroimage 199, 114–126.

Clarke, A., Taylor, K.I., Tyler, L.K., 2011. The evolution of meaning: spatio-temporal dynamics of visual object recognition. J. Cogn. Neurosci. 23, 1887–1899.

Colclough, G.L., Brookes, M.J., Smith, S.M., Woolrich, M.W., 2015. A symmetric multivariate leakage correction for MEG connectomes. Neuroimage 117, 439–448.

Cope, T.E., Shtyrov, Y., MacGregor, L.J., Holland, R., Pulvermüller, F., Rowe, J.B., Patterson, K., 2020. Anterior temporal lobe is necessary for efficient lateralised processing of spoken word identity. cortex 126, 107–118.

Crinion, J.T., Lambon Ralph, M.A., Warburton, E.A., Howard, D., Wise, R.J.S., 2003. Temporal lobe regions engaged during normal speech comprehension. Brain 126, 1193–1201.

Dhond, R.P., Witzel, T., Dale, A.M., Halgren, E., 2007. Spatiotemporal cortical dynamics underlying abstract and concrete word reading. Hum. Brain Mapp. 28, 355–362.

Duncan, J., 2010. The multiple-demand (MD) system of the primate brain: mental programs for intelligent behaviour. Trends Cogn. Sci. 14, 172–179.

Embleton, K. V, Lambon Ralph, M.A., Parker, G.J., 2006. A combined distortion corrected protocol for diffusion weighted tractography and fMRI, in: Proc. Intl. Soc. Mag. Reson. Med. p. 1070.

Engemann, D.A., Gramfort, A., 2015. Automated model selection in covariance estimation and spatial whitening of MEG and EEG signals. Neuroimage 108, 328–342.

Evans, G.A.L., Ralph, M.A.L., Woollams, A.M., 2012. What’s in a word? A parametric study of semantic influences on visual word recognition. Psychon. Bull. Rev. 19, 325–331.

Farahibozorg, S.-R., 2018. Uncovering Dynamic Semantic Networks in the Brain Using Novel Approaches for EEG/MEG Connectome Reconstruction. University of Cambridge.

Farahibozorg, S.-R., Henson, R.N., Hauk, O., 2018. Adaptive cortical parcellations for source reconstructed EEG/MEG connectomes. Neuroimage 169, 23–45.

Farahibozorg, S.-R., Henson, R.N., Woollams, A.M., Hauk, O., 2019. Distinct roles for the Anterior Temporal Lobe and Angular Gyrus in the spatio-temporal cortical semantic network. bioRxiv 544114.

Flick, G., Oseki, Y., Kaczmarek, A.R., Al Kaabi, M., Marantz, A., Pylkkänen, L., 2018. Building words and phrases in the left temporal lobe. Cortex 106, 213–236.

Fuchs, M., Wagner, M., Köhler, T., Wischmann, H.-A., 1999. Linear and nonlinear current density reconstructions. J. Clin. Neurophysiol. 16, 267–295.

Glasser, M.F., Coalson, T.S., Robinson, E.C., Hacker, C.D., Harwell, J., Yacoub, E., Ugurbil, K., Andersson, J., Beckmann, C.F., Jenkinson, M., 2016. A multi-modal parcellation of human cerebral cortex. Nature 536, 171–178.

Gramfort, A., Luessi, M., Larson, E., Engemann, D.A., Strohmeier, D., Brodbeck, C., Goj, R., Jas, M., Brooks, T., Parkkonen, L., 2013. MEG and EEG data analysis with MNE-Python. Front. Neurosci. 7, 267.

Gramfort, A., Luessi, M., Larson, E., Engemann, D.A., Strohmeier, D., Brodbeck, C., Parkkonen, L., Hämäläinen, M.S., 2014. MNE software for processing MEG and EEG data. Neuroimage 86, 446–460.

Hämäläinen, M.S., Ilmoniemi, R.J., 1994. Interpreting magnetic fields of the brain: minimum norm estimates. Med. Biol. Eng. Comput. 32, 35–42.

Hanslmayr, S., Staudigl, T., Fellner, M.-C., 2012. Oscillatory power decreases and long-term memory: the information via desynchronization hypothesis. Front. Hum. Neurosci. 6, 74.

Hauk, O., 2016. Only time will tell–why temporal information is essential for our neuroscientific understanding of semantics. Psychon. Bull. Rev. 23, 1072–1079.

Hauk, O., 2004. Keep it simple: a case for using classical minimum norm estimation in the analysis of EEG and MEG data. Neuroimage 21, 1612–1621.

Hauk, O., Coutout, C., Holden, A., Chen, Y., 2012. The time-course of single-word reading: evidence from fast behavioral and brain responses. Neuroimage 60, 1462–1477.

Hauk, O., Stenroos, M., Treder, M., 2019. Towards an objective evaluation of EEG/MEG source estimation methods: The Linear Tool Kit. BioRxiv 672956.

Hauk, O., Wakeman, D.G., Henson, R., 2011. Comparison of noise-normalized minimum norm estimates for MEG analysis using multiple resolution metrics. Neuroimage 54, 1966–1974.

Humphreys, G.F., Hoffman, P., Visser, M., Binney, R.J., Lambon Ralph, M.A., 2015. Establishing task-and modality-dependent dissociations between the semantic and default mode networks. Proc. Natl. Acad. Sci. 112, 7857–7862.

Humphreys, G.F., Lambon Ralph, M.A., 2014. Fusion and fission of cognitive functions in the human parietal cortex. Cereb. Cortex 25, 3547–3560.

Hyvarinen, A., 1999. Fast and robust fixed-point algorithms for independent component analysis. IEEE Trans. Neural Networks 10, 626–634.

Hyvärinen, A., Oja, E., 2000. Independent component analysis: algorithms and applications. Neural networks 13, 411–430.

Jackson, R.L., 2021. The neural correlates of semantic control revisited. Neuroimage 224, 117444.

Jackson, R.L., Hoffman, P., Pobric, G., Lambon Ralph, M.A., 2016. The semantic network at work and rest: differential connectivity of anterior temporal lobe subregions. J. Neurosci. 36, 1490– 1501.

Jefferies, E., 2013. The neural basis of semantic cognition: converging evidence from neuropsychology, neuroimaging and TMS. Cortex 49, 611–625.

Jefferies, E., Lambon Ralph, M.A., 2006. Semantic impairment in stroke aphasia versus semantic dementia: a case-series comparison. Brain 129, 2132–2147.

Jung, J., Ralph, M.A.L., 2019. Enhancing vs. inhibiting semantic performance with repetitive transcranial magnetic stimulation over the anterior temporal lobe: frequency-and task-specific effects. bioRxiv.

Kietzmann, T.C., Spoerer, C.J., Sörensen, L.K.A., Cichy, R.M., Hauk, O., Kriegeskorte, N., 2019. Recurrence is required to capture the representational dynamics of the human visual system. Proc. Natl. Acad. Sci. 116, 21854–21863.

Kuhnke, P., Kiefer, M., Hartwigsen, G., 2021. Task-Dependent Functional and Effective Connectivity during Conceptual Processing. Cereb. Cortex.

Kuhnke, P., Kiefer, M., Hartwigsen, G., 2020. Task-dependent recruitment of modality-specific and multimodal regions during conceptual processing. Cereb. Cortex 30, 3938–3959.

Kutas, M., Federmeier, K.D., 2011. Thirty years and counting: finding meaning in the N400 component of the event-related brain potential (ERP). Annu. Rev. Psychol. 62, 621–647.

Lachaux, J., Rodriguez, E., Martinerie, J., Varela, F.J., 1999. Measuring phase synchrony in brain signals. Hum. Brain Mapp. 8, 194–208.

Lambon Ralph, M.A., Jefferies, E., Patterson, K., Rogers, T.T., 2016. The neural and computational bases of semantic cognition. Nat. Rev. Neurosci. 18, 42–55. https://doi.org/10.1038/nrn.2016.150

Lambon Ralph, M.A., Sage, K., Jones, R.W., Mayberry, E.J., 2010. Coherent concepts are computed in the anterior temporal lobes. Proc. Natl. Acad. Sci. 107, 2717–2722.

Lamme, V.A.F., Roelfsema, P.R., 2000. The distinct modes of vision offered by feedforward and recurrent processing. Trends Neurosci. 23, 571–579.

Lau, E.F., Gramfort, A., Hämäläinen, M.S., Kuperberg, G.R., 2013. Automatic semantic facilitation in anterior temporal cortex revealed through multimodal neuroimaging. J. Neurosci. 33, 17174–17181.

Lau, E.F., Phillips, C., Poeppel, D., 2008. A cortical network for semantics:(de) constructing the N400. Nat. Rev. Neurosci. 9, 920–933.

Lewis, A.G., Bastiaansen, M., 2015. A predictive coding framework for rapid neural dynamics during sentence-level language comprehension. Cortex 68, 155–168.

Lewis, G.A., Poeppel, D., Murphy, G.L., 2015. The neural bases of taxonomic and thematic conceptual relations: An MEG study. Neuropsychologia 68, 176–189.

Liu, A.K., Dale, A.M., Belliveau, J.W., 2002. Monte Carlo simulation studies of EEG and MEG localization accuracy. Hum. Brain Mapp. 16, 47–62.

Marinkovic, K., Dhond, R.P., Dale, A.M., Glessner, M., Carr, V., Halgren, E., 2003. Spatiotemporal dynamics of modality-specific and supramodal word processing. Neuron 38, 487–497.

Marinkovic, K., Rosen, B.Q., Cox, B., Hagler Jr, D.J., 2014. Spatio-temporal processing of words and nonwords: Hemispheric laterality and acute alcohol intoxication. Brain Res. 1558, 18– 32.

Maris, E., Oostenveld, R., 2007. Nonparametric statistical testing of EEG-and MEG-data. J. Neurosci. Methods 164, 177–190.

Martin, A., 2016. GRAPES—Grounding representations in action, perception, and emotion systems: How object properties and categories are represented in the human brain. Psychon. Bull. Rev. 23, 979–990.

Medler, D.A., Binder, J.R., 2005. MCWord: An On-Line Orthographic Database of the English Language. [WWW Document].

Mion, M., Patterson, K., Acosta-Cabronero, J., Pengas, G., Izquierdo-Garcia, D., Hong, Y.T., Fryer, T.D., Williams, G.B., Hodges, J.R., Nestor, P.J., 2010. What the left and right anterior fusiform gyri tell us about semantic memory. Brain 133, 3256–3268.

Molins, A., Stufflebeam, S.M., Brown, E.N., Hämäläinen, M.S., 2008. Quantification of the benefit from integrating MEG and EEG data in minimum ℓ2-norm estimation. Neuroimage 42, 1069–1077.

Mollo, G., Cornelissen, P.L., Millman, R.E., Ellis, A.W., Jefferies, E., 2017. Oscillatory dynamics supporting semantic cognition: MEG evidence for the contribution of the anterior temporal lobe hub and modality-specific spokes. PLoS One 12, e0169269.

Mummery, C.J., Patterson, K., Price, C.J., Ashburner, J., Frackowiak, R.S.J., Hodges, J.R., 2000. A voxel-based morphometry study of semantic dementia: relationship between temporal lobe atrophy and semantic memory. Ann. Neurol. 47, 36–45.

Mur, M., Ruff, D.A., Bodurka, J., De Weerd, P., Bandettini, P.A., Kriegeskorte, N., 2012. Categorical, yet graded–single-image activation profiles of human category-selective cortical regions. J. Neurosci. 32, 8649–8662.

Nestor, P.J., Fryer, T.D., Hodges, J.R., 2006. Declarative memory impairments in Alzheimer’s disease and semantic dementia. Neuroimage 30, 1010–1020.

Nolte, G., Bai, O., Wheaton, L., Mari, Z., Vorbach, S., Hallett, M., 2004. Identifying true brain interaction from EEG data using the imaginary part of coherency. Clin. Neurophysiol. 115, 2292–2307.

Noonan, K.A., Jefferies, E., Visser, M., Lambon Ralph, M.A., 2013. Going beyond inferior prefrontal involvement in semantic control: evidence for the additional contribution of dorsal angular gyrus and posterior middle temporal cortex. J. Cogn. Neurosci. 25, 1824–1850.

Olson, I.R., Plotzker, A., Ezzyat, Y., 2007. The enigmatic temporal pole: a review of findings on social and emotional processing. Brain 130, 1718–1731.

Palva, J.M., Wang, S.H., Palva, S., Zhigalov, A., Monto, S., Brookes, M.J., Schoffelen, J.-M., Jerbi, K., 2018. Ghost interactions in MEG/EEG source space: A note of caution on inter-areal coupling measures. Neuroimage 173, 632–643.

Patterson, K., Nestor, P.J., Rogers, T.T., 2007. Where do you know what you know? The representation of semantic knowledge in the human brain. Nat. Rev. Neurosci. 8, 976.

Patterson, K., Ralph, M.A.L., Jefferies, E., Woollams, A., Jones, R., Hodges, J.R., Rogers, T.T., 2006. “Presemantic” cognition in semantic dementia: six deficits in search of an explanation. J. Cogn. Neurosci. 18, 169–183.

Pobric, G., Jefferies, E., Lambon Ralph, M.A., 2007. Anterior temporal lobes mediate semantic representation: mimicking semantic dementia by using rTMS in normal participants. Proc. Natl. Acad. Sci. 104, 20137–20141.

Pulvermüller, F., Shtyrov, Y., Hauk, O., 2009. Understanding in an instant: neurophysiological evidence for mechanistic language circuits in the brain. Brain Lang. 110, 81–94.

Rice, G.E., Hoffman, P., Ralph, M.A.L., 2015a. Graded specialization within and between the anterior temporal lobes. Ann. N. Y. Acad. Sci. 1359, 84.

Rice, G.E., Lambon Ralph, M.A., Hoffman, P., 2015b. The roles of left versus right anterior temporal lobes in conceptual knowledge: an ALE meta-analysis of 97 functional neuroimaging studies. Cereb. Cortex 25, 4374–4391.

Rogers, T.T., Hocking, J., Noppeney, U.T.A., Mechelli, A., Gorno-Tempini, M.L., Patterson, K., Price, C.J., 2006. Anterior temporal cortex and semantic memory: reconciling findings from neuropsychology and functional imaging. Cogn. Affect. Behav. Neurosci. 6, 201–213.

Rogers, T.T., Lambon Ralph, M.A., Matthew, A., Garrard, P., Bozeat, S., McClelland, J.L., Hodges, J.R., Patterson, K., 2004. Structure and deterioration of semantic memory: a neuropsychological and computational investigation. Psychol. Rev. 111, 205.

Schoffelen, J.-M., Hultén, A., Lam, N., Marquand, A.F., Uddén, J., Hagoort, P., 2017. Frequency-specific directed interactions in the human brain network for language. Proc. Natl. Acad. Sci. 114, 8083–8088.

Siegel, M., Donner, T.H., Engel, A.K., 2012. Spectral fingerprints of large-scale neuronal interactions. Nat. Rev. Neurosci. 13, 121–134.

Strijkers, K., Bertrand, D., Grainger, J., 2015. Seeing the same words differently: The time course of automaticity and top–down intention in reading. J. Cogn. Neurosci. 27, 1542–1551.

Taulu, S., Kajola, M., 2005. Presentation of electromagnetic multichannel data: the signal space separation method. J. Appl. Phys. 97, 124905.

Tranel, D., Grabowski, T.J., Lyon, J., Damasio, H., 2005. Naming the same entities from visual or from auditory stimulation engages similar regions of left inferotemporal cortices. J. Cogn. Neurosci. 17, 1293–1305.

Visser, M., Embleton, K. V, Jefferies, E., Parker, G.J., Lambon Ralph, M.A., 2010. The inferior, anterior temporal lobes and semantic memory clarified: novel evidence from distortion-corrected fMRI. Neuropsychologia 48, 1689–1696.

Visser, M., Jefferies, E., Embleton, K. V, Lambon Ralph, M.A., 2012. Both the middle temporal gyrus and the ventral anterior temporal area are crucial for multimodal semantic processing: distortion-corrected fMRI evidence for a double gradient of information convergence in the temporal lobes. J. Cogn. Neurosci. 24, 1766–1778.

Wens, V., Marty, B., Mary, A., Bourguignon, M., Op de Beeck, M., Goldman, S., Van Bogaert, P., Peigneux, P., De Tiège, X., 2015. A geometric correction scheme for spatial leakage effects in MEG/EEG seed-based functional connectivity mapping. Hum. Brain Mapp. 36, 4604–4621.

Westerlund, M., Pylkkänen, L., 2014. The role of the left anterior temporal lobe in semantic composition vs. semantic memory. Neuropsychologia 57, 59–70.

Williams, A., Reddigari, S., Pylkkänen, L., 2017. Early sensitivity of left perisylvian cortex to relationality in nouns and verbs. Neuropsychologia 100, 131–143.

Williams, N., Arnulfo, G., Wang, S.H., Nobili, L., Palva, S., Palva, J.M., 2019. Comparison of methods to identify modules in noisy or incomplete brain networks. Brain Connect. 9, 128– 143.

Woodhead, Z.V.J., Barnes, G.R., Penny, W., Moran, R., Teki, S., Price, C.J., Leff, A.P., 2014. Reading frosnt to back: MEG evidence for early feedback effects during word recognition. Cereb. Cortex 24, 817–825.

